# Long-term natural selection affects patterns of neutral divergence on the X chromosome more than the autosomes

**DOI:** 10.1101/023234

**Authors:** Pooja Narang, Melissa A. Wilson Sayres

**Affiliations:** School of Life Sciences, Arizona State University, Tempe, Arizona, 85281 USA; Center for Evolution and Medicine, The Biodesign Institute, Arizona State University, Tempe, Arizona, 85281 USA

**Keywords:** X chromosome, background selection, male mutation bias, natural selection, divergence

## Abstract

Natural selection reduces neutral population genetic diversity near coding regions of the genome because recombination has not had time to unlink selected alleles from nearby neutral regions. For ten sub-species of great apes, including human, we show that long-term selection affects estimates of divergence on the X differently from the autosomes. Divergence increases with increasing distance from genes on both the X chromosome and autosomes, but increases faster on the X chromosome than autosomes, resulting in increasing ratios of X/A divergence in putatively neutral regions. Similarly, divergence is reduced more on the X chromosome in neutral regions near conserved regulatory elements than on the autosomes. Consequently estimates of male mutation bias, which rely on comparing neutral divergence between the X and autosomes, are twice as high in neutral regions near genes versus far from genes. Our results suggest filters for putatively neutral genomic regions differ between the X and autosomes.

## Introduction

Natural selection on regions of the genome that affect traits can also affect the evolution of nearby neutral regions that are less likely to be separated from the selected allele by recombination. For example, levels of diversity are reduced in coding genes and in the regions around genes, likely because purifying selection removes harmful alleles and nearby neutral sites are affected by background selection (Charlesworth 2012), or because positive selection increases the frequency of beneficial alleles and nearby neutral sites are affected by hitchhiking (Smith and Haigh 1974). There has been substantial effort to understand how natural selection affects genetic variation within populations, genetic diversity, at putatively neutral sites (Nielsen et al. 2007; Akey 2009; Lohmueller et al. 2011; Wilson Sayres et al. 2014). In humans, genetic diversity is reduced near genes and other conserved sequences (Hammer et al. 2004; Hammer et al. 2008; Hammer et al. 2010; Gottipati et al. 2011; Arbiza et al. 2014), consistent with selection reducing genetic variation at and near selected regions. Further, selection affects diversity differently on the X chromosome and the autosomes; genetic diversity on chromosome X increases faster with increasing distance from genes than it does on the autosomes across human populations (Hammer et al. 2010; Gottipati et al. 2011; Prado-Martinez et al. 2013; Arbiza et al. 2014). Recombination over long evolutionary time is expected to unlink selected sites from linked neutral loci, but it has been suggested that selection may affect the accumulation of substitutions and affect estimates of species divergence in neutral regions near and far from genes on the autosomes (McVicker et al. 2009; Lohmueller et al. 2011). It is not known whether this long-term effect differs between the sex chromosomes and the autosomes.

If natural selection affects substitution rates in neutrally evolving regions differently on the X and autosomes, it will have a tremendous impact on estimates of sex-biased mutation rate differences. In mammals, the accumulation of mutations occurs at different rates in the male and female germlines, being higher in males than in females, a male mutation bias, due to more germline cell divisions in males versus females (Drake et al. 1998). Substitution rates in neutral regions on the X and autosomes are used as a proxy for mutation rate to estimate the ratio of the mutation rate in males to the mutation rate in females (α), because they spend different amounts of time in the male and female germline (Miyata et al. 1987; Ellegren 2007; Wilson Sayres and Makova 2011). Genome-wide substitution rate comparisons are less prone to locus specific variation, and more likely to reflect the true male mutation bias (Li et al. 2002; Makova et al. 2004; Taylor et al. 2006), perhaps not only because of variation in rates of evolution across the genome, but because of differential effects of linked selection on the X and autosomes.

Here, we investigate patterns of divergence with increasing distance from genes on the X chromosome and the autosomes in ten sub-species of great apes (Prado-Martinez et al. 2013). We find that genetic divergence typically increases with increasing genetic distance from genes, and notably that this increase is faster on the X chromosome than the autosomes. That natural selection appears to affect divergence differently on the X and autosomes results in a non-constant ratio of X/A divergence with distance from genes. Consequently, measurements of male mutation bias also vary with distance from genes, with estimates near genes almost twice as high as those far from genes. Our results suggest that natural selection shapes patterns of divergence in neutral genomic regions across the great apes, with unique effects on the X and autosomes that can significantly impact estimates of evolutionary processes that compare substitution rates between these regions of the genome.

## Results & Discussion

We find that divergence increase as the distance from the genes increases on both the X and the autosomes (Figure 1), suggesting that selection acts on linked neutral regions even over long evolutionary time. Interestingly, similar to patterns of diversity within humans (Hammer et al. 2010; Gottipati et al. 2011; Arbiza et al. 2014), we also observe that divergence on the X chromosome increases faster than the autosomes with increasing distance from genes (Figure 1), consistent with selection on linked neutral sites having a stronger effect on the X chromosome than on the autosomes across the great apes. (Hudson and Kaplan 1995; Charlesworth 1996; Orr and Betancourt 2001; Vicoso and Charlesworth 2006; Ávila et al. 2015).

If selection is affecting estimates of divergence near and far from genes differently on the X chromosome and the autosomes, as we observe, then the ratio of divergence on the X versus the autosomes will vary depending on which regions are analyzed. We observe that the ratio of X/A divergence increases with increasing distance from genes, driven by a faster increase in the rate of divergence on the X chromosome relative to the autosomes (Figure S1). However, because of the different effective population sizes of the chromosomes, estimates of divergence may be inflated differently on the X and autosomes by variable levels of ancestral polymorphism for each chromosome type (Li et al. 2002; Makova and Li 2002). This correction for ancient diversity is expected to reduce the divergence on both X and autosomes, but the reduction in divergence on X is expected to be less due to a smaller effective population size and lower expected ancestral polymorphism (Li et al. 2002). To account for this, across all great apes, we compute ratios of X/A divergence corrected for ancestral polymorphism (Materials and Methods), and find that corrected X/A divergence ratios are indeed higher than the uncorrected ratios (Figure S1)

**Figure 1.**
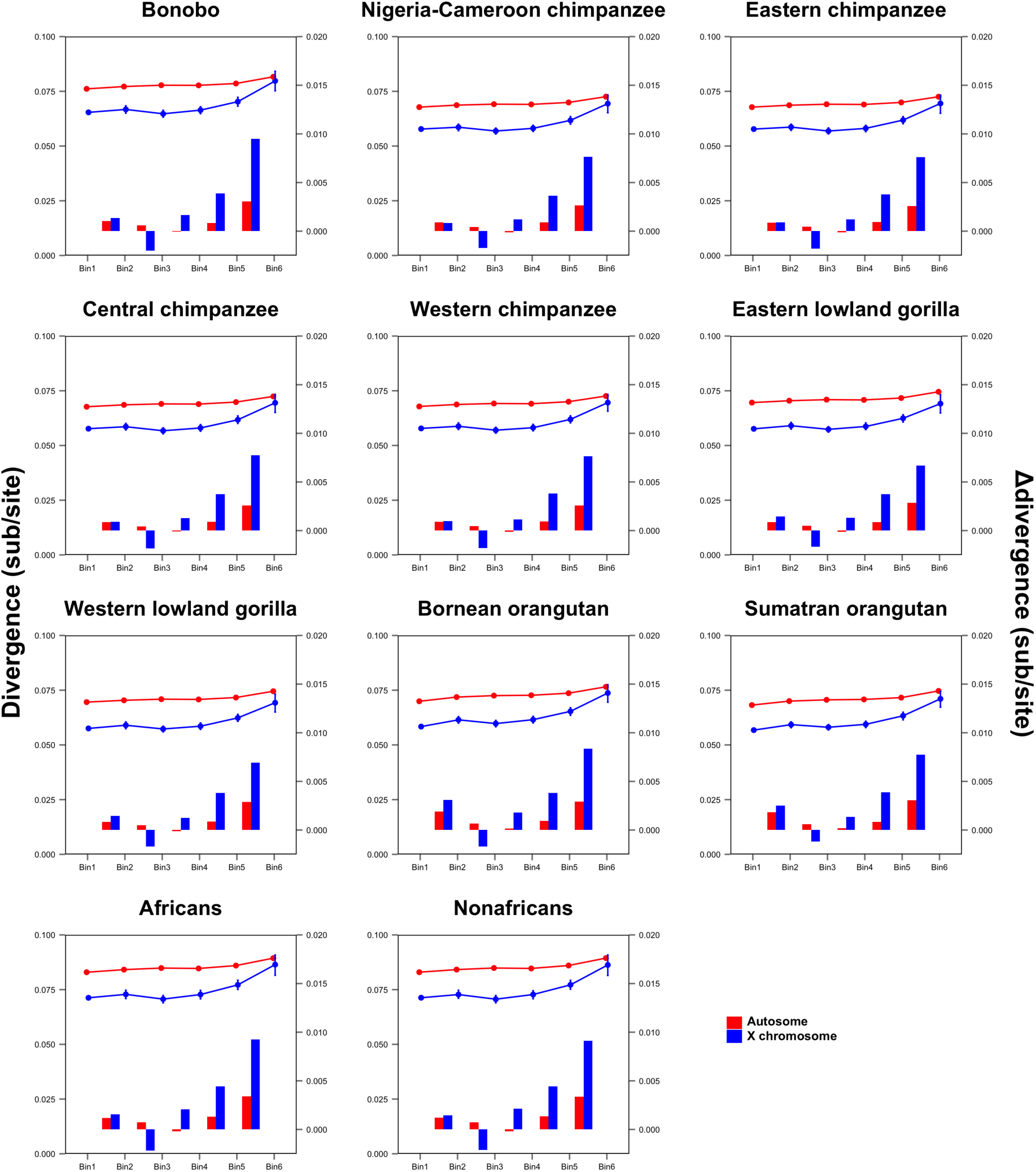
Divergence across great apes in neutral regions with increasing distance from genes for autosomes and X chromosome. Divergence increases with distance from genes faster on the X chromosome (blue) than on the autosomes (red) for all great ape subspecies, plotted as lines. 95% confidence intervals calculated using bootstrap procedure are plotted. The difference between divergences of two consecutive bins is plotted as bars between each pair of bins. (Bin1=0-0.05cM, Bin2=0.05-0.1cM, Bin3=0.1-0.2cM, Bin4=0.2-0.4cM, Bin5=0.4-0.8cM, Bin6=0.8-2.0cM).

Male mutation bias, the ratio of the mutation rate in males to females (α), is estimated from genetic divergence data on the X and autosomes, assuming they spend different amounts of time in the male and female germline (Miyata et al. 1987; Li et al. 2002; Makova and Li 2002; Makova et al. 2004; Taylor et al. 2006; Wilson Sayres and Makova 2011). However, if natural selection affects divergence patterns on the X and autosomes differently near and far from genes, then genomic regions analyzed will greatly affect estimates of α from substitution rates. Using both the corrected and uncorrected X/A divergence ratios, we compute α with increasing distance from genes. For all great apes we observe higher magnitudes of α in genomic regions near genes, and lower estimates far from genes (Figure S2). Estimates of male mutation bias near genes are twice as high as estimates furthest from the genes (Figure 2A). The absolute estimates of α are very similar for all the subspecies due to averaging the divergence estimates over long evolutionary times from each modern species to the most recent common ancestor of all great apes (Figure 2B; Materials and Methods). The low estimate of α, far from genes, as averaged over 10.5 million years of evolution suggests that in the common ancestor of the great apes, and for much of great ape evolutionary history, there was not much difference in the mutation rate between males and females. Further, even averaged over long evolutionary time the variation in estimates of α near and far from genes suggests the importance of considering the long-term differential effects of selection on rates of divergence across genomic regions.

**Figure 2.**
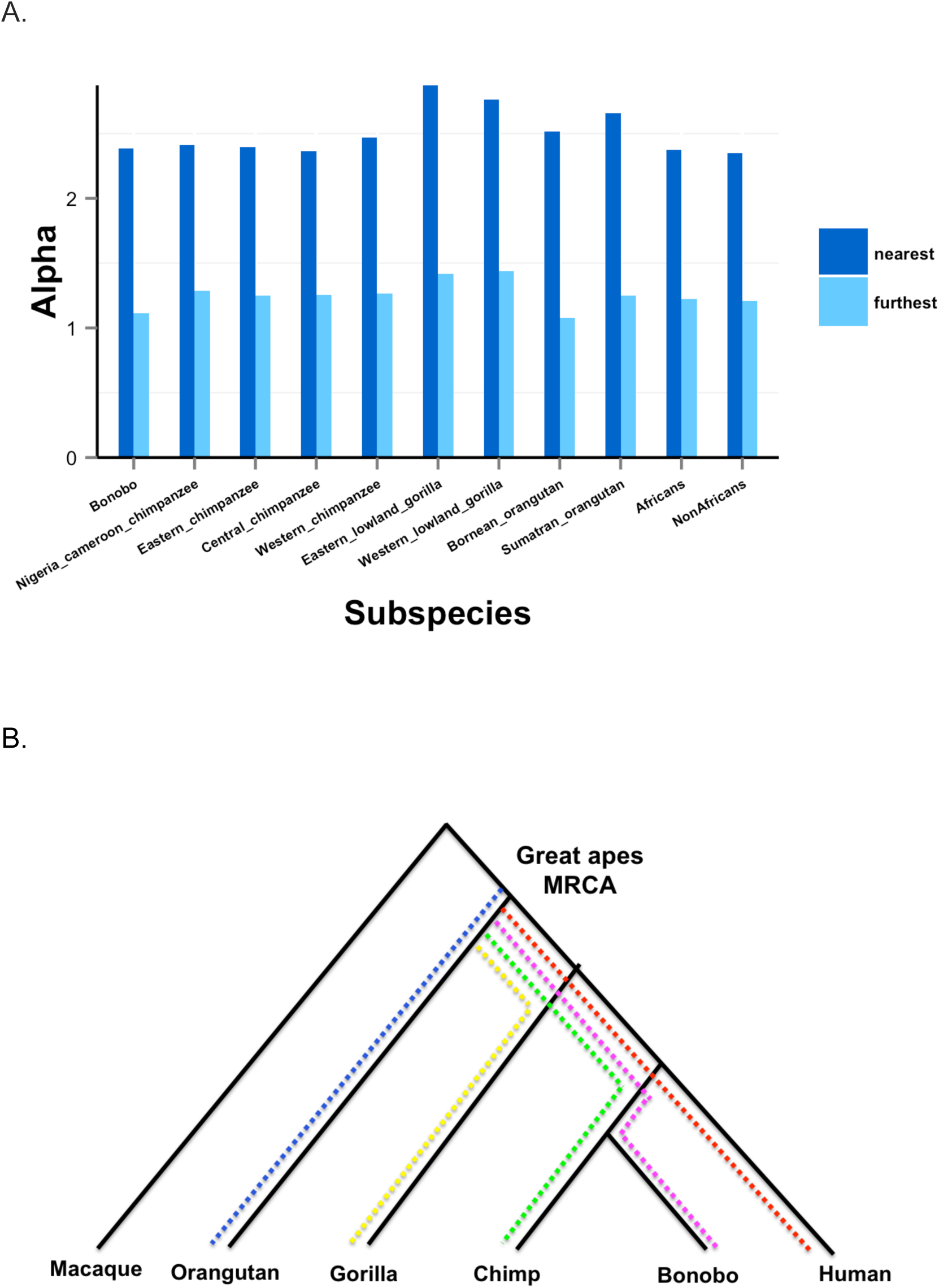
Male mutation bias computed along each ape branch to the Great Ape Most Recent Common Ancestor, MRCA. A. Male mutation bias estimates, computed as α, nearest and furthest from the genes. B. Divergence estimates are calculated for each sub-species to the most recent common ancestor, MRCA of the great apes.

### Analysis to explain variation in the 0.1-0.2 cM genic region

Interestingly, while divergence increases monotonically on the autosomes, this is not true for the X chromosome (Figure 1). We observe that divergence initially increases with increasing distance from genes, then observe a dip in divergence in the bin [0.1-0.2] cM from genes on the X chromosome (Figure 1). Differences in divergence between adjacent bins suggest that this dip is isolated, and driven by a particular genomic feature, because divergence on the X chromosome increases consistently with distance from genes after this bin (Figure 1). We also observe the lowest X/A ratio and the highest estimates of male mutation bias (α _corrected_ & α _uncorrected_) in the [0.1-0.2] cM from genes bin (Figure S2), driven by the low divergence on the X relative to the autosomes in this bin.

To investigate the cause in the reduction in divergence on the X chromosome [0.1-0.2] cM from genes, we examined if this region is associated with any conserved or putative regulatory elements. We calculated the frequency of CpG elements, repeat elements, simple repeats and phastCons in six bins with increasing physical distance from genes both for autosomes and the X chromosome (Figure S3; Materials and Methods). Generally, the frequency of each element decreases as the distance from the genes increases. We do not observe a higher occurrence of CpG elements, repetitive elements, or simple repetitive elements in the [0.1-0.2] Mbp from genes bin for either the autosomes or X-chromosome (Figure S3A,B). However, we do observe an increase in the frequency of phastCons elements on the X chromosome in this region [0.1-0.2] Mbp, with only a marginal increase in the frequency of phastCons on the autosomes in this bin (Figure S3C,D). Thus, we hypothesize that natural selection acting on conserved non-coding elements reduces divergence on the X chromosome more than the autosomes.

To test the effect of phastCons elements on reducing divergence, we compute branch-specific divergence with and without these regions for the X and the autosomes (Materials and Methods). The ratio of X/A divergence (both uncorrected and corrected for ancestral polymorphism) increases with the removal of phastCons elements (Figure 3), indicating that phastCons, which are subject to long-term purifying selection that reduces divergence (POLLARD ET AL. 2010), contribute to reducing divergence more on the X chromosome than the autosomes. We see a further increase in the X/A divergence ratio when an additional 10kb flanking the phastCons are also removed, again driven by an increase in divergence on the X chromosome (Figure 3). These results suggest that selection to preserve phastCons elements also affects variation at linked neutral sites, and that this linked selection (likely background selection) is stronger on the X than the autosomes.

**Figure 3.**
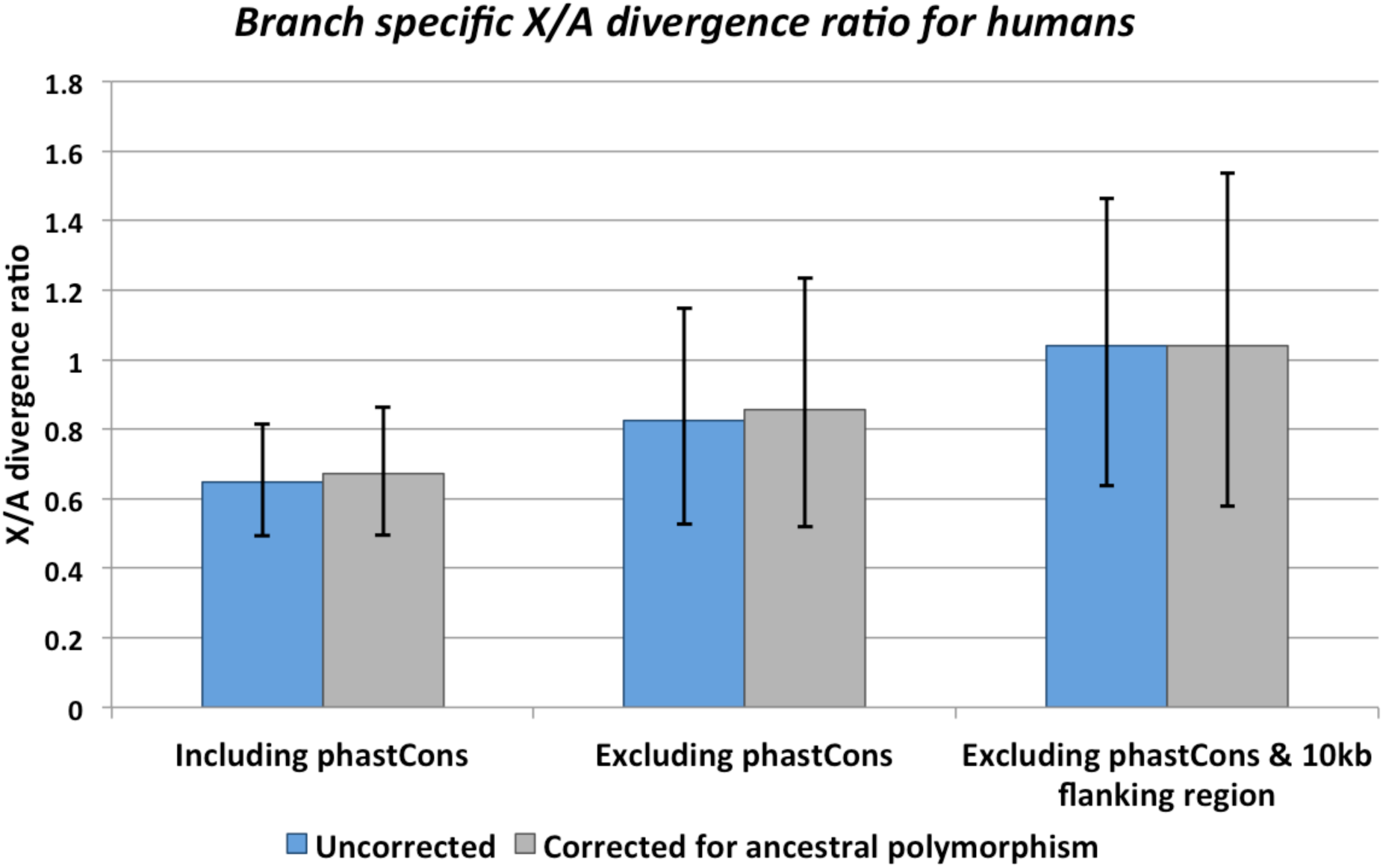
Divergence on the X chromosome and autosomes including and excluding phastCons and nearby neutral regions. X to autosomal divergence ratio calculated for three cases: (i) neutral regions including phastCons; (ii) neutral regions without phastCons; and (iii) neutral regions without phastCons and without 10kb flanking regions around phastCons.

## Conclusions

We studied X and autosomal divergence and the ratio of X/A divergence for ten subspecies of the great apes. The our results show that divergence increases with increasing distance from genes faster on the X chromosome than the autosomes, signifying the role of selection in differentially shaping patterns of neutral divergence on the X and autosomes. We further show that the differential effects of linked selection on neutral sites on the X chromosome and autosome will affect analyses that compare the rates of evolution on these two regions. In particular, estimates of male mutation bias close to the genes were twice as high as estimates farthest from genes, driven by a faster divergence with distance from genes on the X chromosome. These results shed light on how selection shapes patterns of divergence across species, and between genomic regions.

## Materials and Methods

### Sequence data

We analyzed data from whole genome sequences of 77 individuals from ten great ape sub-species (*Homo sapiens*, *Pan paniscus, Pan troglodytes ellioti, Pan troglodytes schweinfurthii, Pan troglodytes troglodytes, Pan troglodytes verus, Gorilla beringei graueri, Gorilla gorilla gorilla, Pongo pygmaeus, and Pongo abelii*) (Prado-Martinez et al. 2013). Nucleotide diversity and divergence values, calculated for callable bases in 20kbp windows were retrieved from the original work and divided into six bins of increasing genetic distances from the nearest genes (in centiMorgans) as follows: ([0-0.05], [0.05-0.1], [0.1-0.2], [0.2-0.4], [0.4-0.8], [0.8-2.0]). Divergence for each great ape subspecies was computed along the branch to the common ancestor of all great apes (Figure 2), meaning the branches for each sub-species cover the same amount of evolutionary time (Prado-Martinez et al. 2013).

### Genomic coordinates across the great ape population data

For comparison across great ape subspecies populations we extracted positions of coding RefSeq genes, CpG islands, repetitive elements, simple repeats, and phastCons elements from the UCSC Genome Browser (Fujita et al. 2011) for the hg18 build of the human reference genome because the great apes diversity and divergence values were estimated using the same build (Prado-Martinez et al. 2013). Distance of the conserved and regulatory elements to the RefSeq gene positions is calculated using the closest-features tool from BEDOPS program (Neph et al. 2012). The frequency of each element is computed in bins using physical distance (in base pairs) from the genes is calculated here, considering an approximate conversion of 1cM ∼ 1Mbp (Lodish et al. 2004).

### Estimating divergence across reference genomes

We used the Neutral Region Explorer webserver (Arbiza et al. 2012) to extract putatively neutral regions for the hg19 human reference genome, masking the genome for genic regions (known genes, gene bounds, spliced ESTs), duplicated regions (segmental duplications, copy number variants, self-chain regions), repetitive regions (simple repeats, repetitive elements), phastCons (44wayPlacental; (Pollard et al. 2010)), and 10kb flanking regions around genes and around phastCons. We required filtered regions to have at least 500bp to be included in further analysis. We extracted three sets: (i) regions far from genes (10kb away from genes) including phastCons; (ii) regions far from genes (i.e in (i) above) and removing phastCons; and, (iii) regions far from genes (i.e in (i) above) and removing both phastCons and 10kb flanking regions around phastCons. For each filtered region we extracted a five-way multiple sequence alignment including the reference genomes of human (hg19), chimp (panTro4), gorilla (gorGor3), orangutan (ponAbe2), and macaque (rheMac3) using the Galaxy interface (Goecks et al. 2010). Branch-specific substitution rates were calculated using using PhyML (Guindon et al. 2010). Diversity estimates for CEU populations, obtained from the NRE webserver for the same intervals descried above (Arbiza et al. 2012), were used to correct for ancestral diversity when estimating the human branch-specific divergence. We used the seqboot tool from phylip (Felsenstein 2005) to generate 1000 alignment data sets for bootstrap analysis and to generate 95% confidence intervals.

### Computing male mutation bias

Within each ape subspecies we corrected for different levels of ancestral polymorphism on the X and autosomes separately by subtracting diversity on each chromosome from the calculated divergence values within each genomic window (Ebersberger et al. 2002; Li et al. 2002; Makova and Li 2002). Both uncorrected and corrected X/A divergence ratios are calculated for each bin of increasing genetic distance from genes for each subspecies. Next we computed uncorrected and corrected male mutation bias (α) for each bin using uncorrected and corrected X/A divergence ratios respectively, according to Miyata’s formula (Miyata et al. 1987):

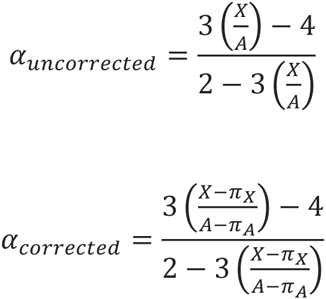

where X and A are X chromosomal and autosomal divergences, respectively, and π_X_ and π_A_ are X chromosomal and autosomal diversities, respectively.

### Statistics

The 95% confidence intervals (CI) are computed using the bootstrap method, where 1000 replicates with replacement are generated from the observed data for each bin (https://github.com/WilsonSayresLab/MMB_apes).

## Acknowledgements

This work was supported by start-up funds from the School of Life Sciences and the Biodesign Institute at Arizona State University to M.A.W.S.. We thank Krishna Veeramah and August Woerner for sharing previously computed data, and for discussions.

